# Do drugs with biliary toxicity cause cholangiocarcinoma?

**DOI:** 10.64898/2026.03.31.715688

**Authors:** Cai Zong, Kyumin Lim, Sierra A. Walker, Rui Dai, Mi Ho Jeong, Reanna George, Jung Hyun Jo, Shams Iqbal, Hyungsoon Im, Ralph Weissleder

**Affiliations:** Center for Systems Biology, Massachusetts General Hospital, Boston, MA 02114, USA; Department of Radiology, Massachusetts General Hospital, Boston, MA 02114, USA; Faculty of Pharmaceutical Sciences, Tokyo University of Science, Tokyo 1258585, Japan; Division of Gastroenterology, Department of Internal Medicine, Yonsei University College of Medicine, Seoul, South Korea; Department of Systems Biology, Harvard Medical School, 200 Longwood Ave, Boston, MA 02115

**Keywords:** cholangiocarcinoma, antibiotic, cancer, drug toxicity

## Abstract

Many commonly used therapeutic drugs cause biliary toxicity, but it is unclear if they are directly responsible for the increasing incidence of cholangiocarcinoma (CCA). We tested experimentally and analyzed through a cohort approach whether drugs, such as the commonly used antibiotic Augmentin, which is a poster-child of biliary toxicity, are causally linked to CCA development. Using sophisticated analytical tools in cholangiocytes, including single extracellular vesicle (EV) analysis, we found no evidence that Augmentin increases the cholangiocyte malignancy marker YAP1 or phospho-YAP1. Furthermore, we analyzed the CCA incidence in our healthcare system and determined it to be 0.0932% (Augmentin group) and 0.0799% (amoxicillin control group). Although the Augmentin group showed a numerically higher CCA incidence, the association did not reach statistical significance (RR = 1.1669, 95% CI 0.6200–2.1961; Fisher’s exact test, P = 0.7493). Similarly, we found no evidence for cholangiocarcinoma development with other commonly used drugs, including chlorpromazine, floxuridine, 5-fluorouracil, flucloxacillin and terbinafine. We conclude that there is no direct causal relationship between clinical Augmentin doses and CCA development.

## Introduction

Cholangiocarcinoma (CCA), cancer of the bile ducts, is a relatively rare malignancy, with an incidence of 0.3–6 cases per 100,000 population per year, with higher rates in Asia. In the United States and worldwide, the incidence and mortality rates of CCA have been steadily increasing over the past few decades, with many plausible associations but no definitive attributable cause(*1*). Despite its low incidence, it poses major clinical challenges because it is often diagnosed at an advanced stage, has a very poor prognosis (5-year survival roughly 7–20%), and has limited effective systemic therapies, with many patients unable to proceed beyond first-line treatment(*2*). The economic burden is substantial: patients frequently require complex care with high healthcare utilization, with U.S. estimates showing treatment costs that easily surpass $250,000/yr with modern targeted agents. Combined with high mortality and increasing incidence, cholangiocarcinoma represents a growing public health concern with significant unmet clinical need and financial impact on healthcare systems.

Screening for cholangiocarcinoma is not recommended in the general population due to its low incidence and lack of highly sensitive and specific screening tools. However, targeted surveillance may be beneficial in high-risk groups, particularly patients with primary sclerosing cholangitis, certain congenital biliary disorders (e.g., choledochal cysts), or chronic biliary inflammation. A much less recognized risk factor is chemical carcinogens and drug-induced cancers. For example, a high incidence of CCA has been reported in Japan among print workers exposed to the organic solvent 1,2-dichloropropane(*3*). Several commonly used drugs have also been associated with adverse effects on cholangiocytes and bile ducts(*4, 4*). These drugs include antibiotics (Augmentin, terbinafine, flucloxacillin)(*5-9*), antineoplastic drugs (floxuridine, 5-fluorouracil)(*10, 11*), immune/anti-inflammatory drugs (Ibuprofen)(*12*), herbal and dietary supplements(*13*), and neurologic drugs (chlorpromazine)(*14*). While there is a clear and well-established correlation between many of these drugs and cholestatic liver injury, it is still unclear whether they cause malignant transformation.

Given the rising incidence of CCA and the frequent use of chemotherapeutics and antibiotics in the US, we asked whether there could be a direct link between certain drugs and CCA development. We initially focused on Augmentin as a commonly used antibiotic associated with biliary toxicity(*15, 16*). Augmentin is a combination antibiotic containing amoxicillin and clavulanic acid that treats a wide range of bacterial infections by both killing bacteria and blocking resistance mechanisms, and is prescribed at ∼30 million times/year in the US alone. We next expanded our analysis to other drugs associated with bile duct injury, including chlorpromazine, floxuridine, 5-fluorouracil, flucloxacillin, and terbinafine. Here, we show no evidence of a causal link between these drugs and cholangiocarcinoma.

## Results

### Augmentin-stressed cholangiocyte model

To evaluate the effects of Augmentin-induced stress on cholangiocytes, we first sought to establish a drug-stress model using the H69 human normal cholangiocyte cell line (**Fig. 1a**). To determine the suitable concentrations of Augmentin stress, H69 cholangiocytes were treated with Augmentin at various concentrations ranging from 0.01 mM to 3.2 mM for 48 h, followed by evaluation of general cytotoxicity. Reported peak plasma concentrations (C_max_) of oral amoxicillin/clavulanate vary by formulation and dose; for example, after an 875/125 mg tablet, the mean amoxicillin C_max_ is approximately 11.6 μg/mL, whereas the clavulanate C_max_ is approximately 2.2 μg/ mL(*17*). Thus, the calculated C_max_ for Augmentin is around 0.032 mM. The cell viability test showed that concentrations below 1 mM did not significantly reduce cell viability (**Fig. 1b, c**), and cytotoxicity only became apparent above 3.2 mM. Accordingly, in this study, we selected 0.32 and 1 mM, which are 10- and 30-fold higher than the reported C_max_ dose, as Augmentin stress doses in our model.

**Fig. 1:**
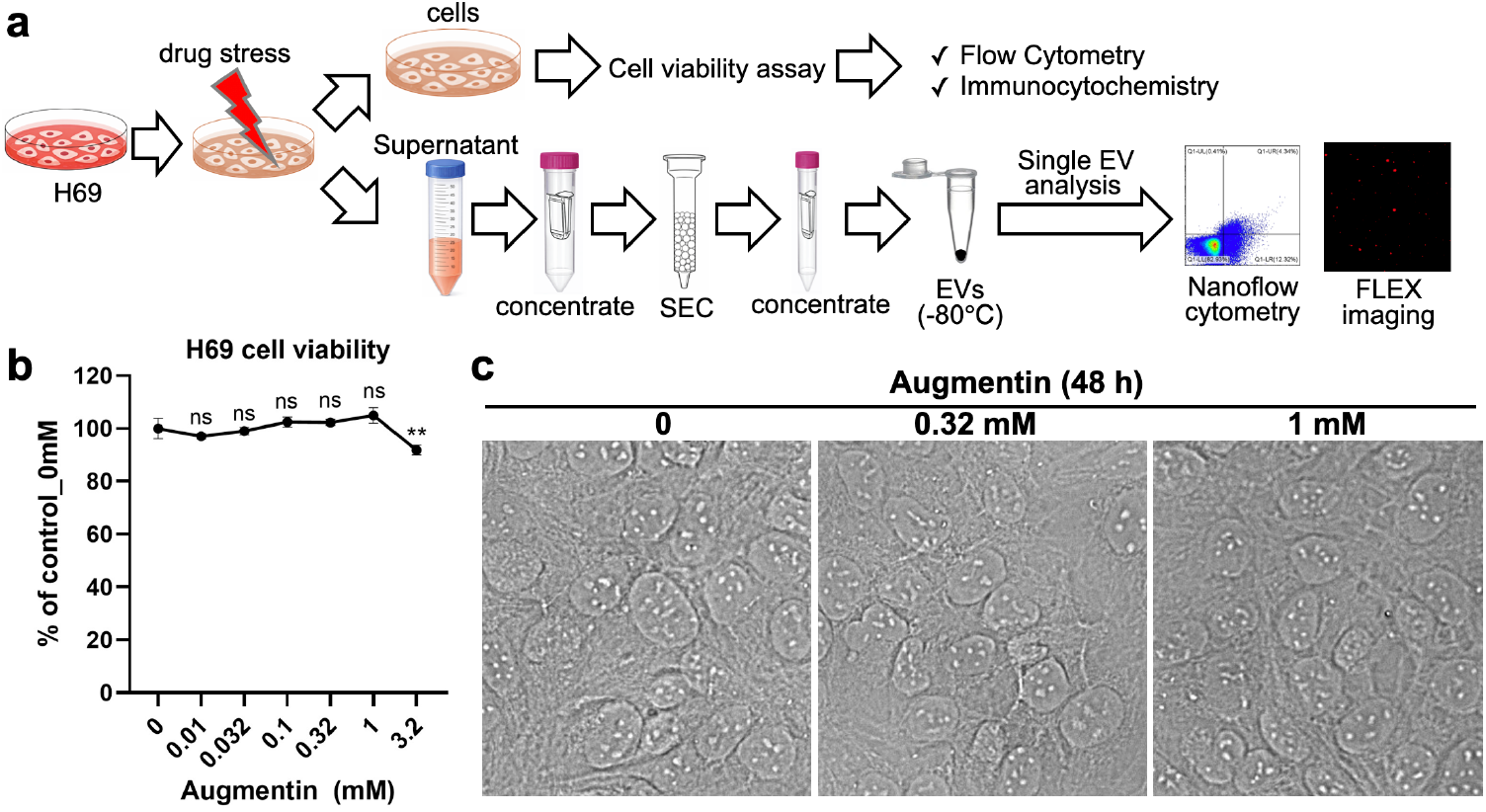
Overview of the established drug-stressed cholangiocyte model and analysis procedures. **(a)** The H69 human normal cholangiocytes were stressed with the targeted drug (e.g., Augmentin) for 48 h. After exposure, the cells were used to evaluate the effects of drug stress on cholangiocyte cell viability and on the expression of selected markers (e.g., the malignancy marker YAP1 and its phosphorylated form, phospho-YAP1) by flow cytometry or immunocytochemistry. In addition, the conditioned medium was collected for isolation of extracellular vesicles (EVs) using size-exclusion chromatography (SEC). The isolated EVs were then subjected to single-EV analyses by nanoflow cytometry or the FLEX platform, including investigation of the effects of drug stress on EV release, EV tetraspanin markers (e.g., CD9/CD63/CD81), and the intra-EV malignancy marker (e.g., YAP1). **(b)** To determine suitable non-lethal drug-stress doses, the effects of Augmentin stress on H69 cholangiocyte cell viability after 48 h of treatment were evaluated using a CCK-8 assay. Up to 1 mM Augmentin-stress for 48 h did not reduce H69 cell viability. Thus, 0.32 and 1 mM, corresponding to 10× and 30× the reported maximum plasma concentration (C_max_), were selected as Augmentin-stress concentrations in our model. **(c)** Microscopic observation confirmed that Augmentin-stress did not significantly affect cholangiocyte morphology at the selected stress doses of 0.32 and 1 mM for 48 h.

### Effects of Augmentin-stress on extracellular vesicle (EV) release and EV tetraspanin markers in H69 cholangiocytes

Analysis of extracellular vesicles (EVs) is emerging as a valuable diagnostic tool in cancer detection, treatment analysis, and toxicity monitoring. As surrogate biomarkers, EVs carry proteins that reflect the molecular status of their originating cells(*18, 19*). In this study, we applied single EV analytical approaches(*18*) in the Augmentin-stressed cholangiocyte model to explore changes in EV markers associated with bile-duct injury and malignancy. We first examined whether Augmentin stress affects the amount of EVs released by cholangiocytes. Increased EV release has been reported as a marker under certain stimuli, for example, lipopolysaccharide (LPS)(*20*) or hypoxia(*21*). EVs were isolated from the conditioned media by size exclusion chromatography after Augmentin treatment for 48 h (**Fig. 1a**). Then, EV counts were measured by nanoflow cytometry, a specialized form of flow cytometry adapted for high-throughput measurement of individual particles at the nanoscale. It first detects particles from machine noise using side-scattering signals, and excludes large aggregates from the analysis (**Fig. 2a**). The results of nanoflow cytometry showed no significant changes in EV counts after Augmentin treatment at 0.32 and 1 mM for 48 h (**Fig. 2b**). Next, we evaluated EV tetraspanin markers, including CD9, CD63, and CD81, as well as their mixture as a pan-CD marker. EVs were universally stained with TFP-555 followed by immunolabeling with antibodies against specific tetraspanin markers for nanoflow cytometry analysis (**Fig. 2c**). Our results showed that among the tested EV tetraspanin markers, CD9-positive EVs were expectedly the most abundant EV subpopulation released from H69 cholangiocytes, followed by the CD63-positive EV subpopulation, whereas the CD81-positive EV subpopulation was negligible. Augmentin stress at 0.32 mM and 1 mM slightly increased CD9, CD63, and pan-CD markers, although the increase was not statistically significant (**Fig. 2d-g**).

**Fig. 2:**
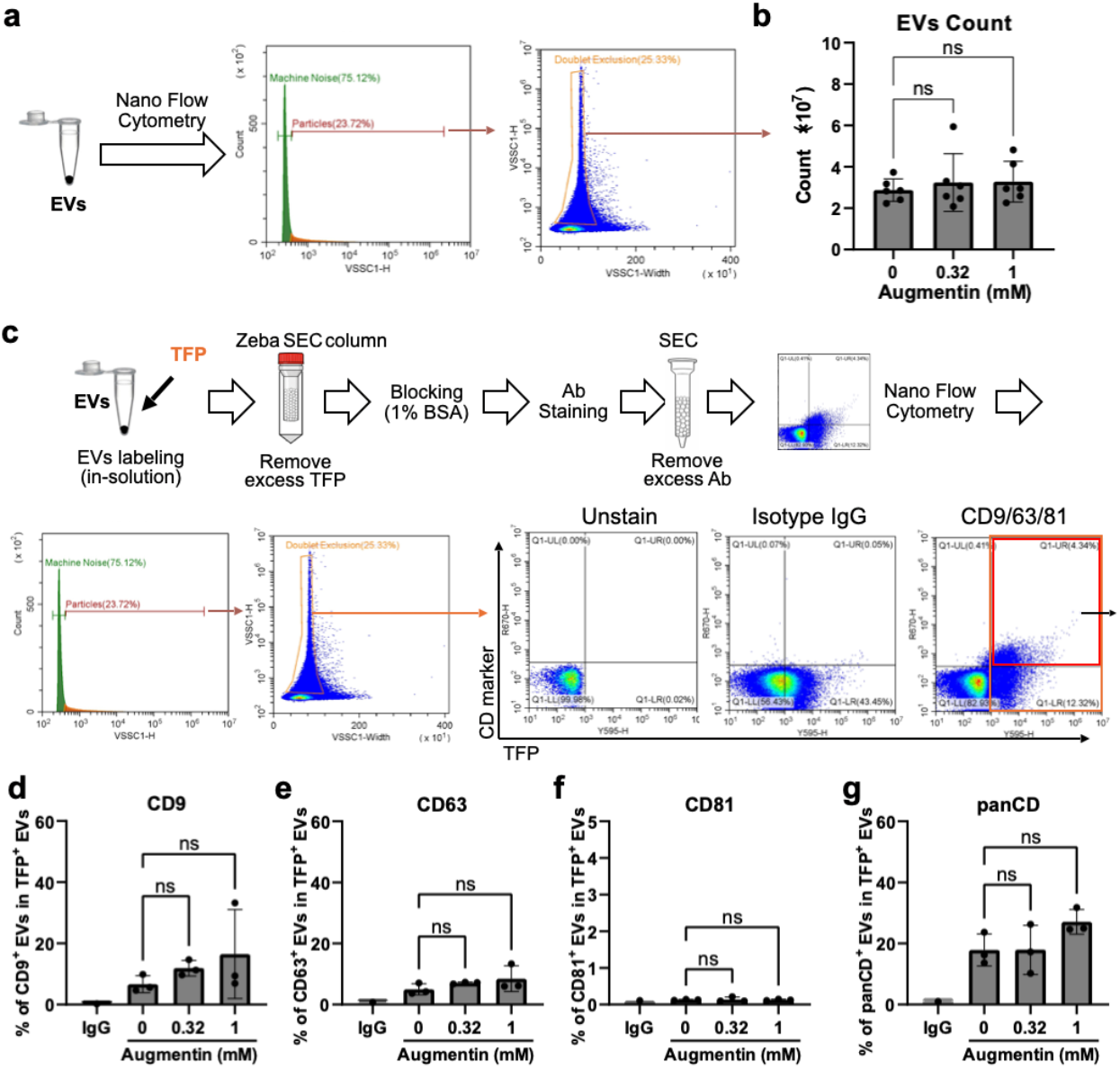
Effects of Augmentin-stress on EV release and EV tetraspanin markers. **(a)** EVs were isolated from the conditioned medium of Augmentin-stressed cholangiocytes and subjected to nanoflow cytometry for EV counts. The gating strategies for the obtained data included separating EV signals from machine noise and further gating to exclude doublets. The number of obtained singlets was calculated as the total EV count of each sample. **(b)** Results of total counts of EVs isolated from cholangiocytes after Augmentin-stress at 0.32 and 1 mM for 48 h. **(c)** Evaluation of EV tetraspanin markers after Augmentin-stress by nanoflow cytometry. Isolated EVs were labelled using pan-EV staining with TFP-555 followed by antibody staining for specific EV tetraspanin markers. Quadrant gating was performed using the TFP-555 signal (X-axis) and marker-647 signal (Y-axis). The percentage of marker-positive EVs among TFP-AF555-positive EVs was calculated as the colocalization ratio of the marker. (**d-g**) Effects of Augmentin-stress on EV tetraspanin markers, including CD9 (d), CD63 (e), CD81 (f), and their mixture as a panCD (g).

### Effects of Augmentin-stress on the cholangiocyte malignancy marker YAP1 in H69 cells and EVs

To further investigate the effects of Augmentin-stress on cholangiocytes, we measured YAP1, a well-established malignancy marker in cholangiocytes(*22*). Hippo/YAP1 signaling is crucially involved in pathophysiology, including development, homeostasis, regeneration, and malignancy in diverse cell types(*23*). In preclinical models, YAP1 activation in cholangiocytes or hepatocytes is sufficient to drive their malignant transformation into CCA(*22*). The results of flow cytometry showed no significant changes in the expression of YAP1 (Fig. 3a-b) and its phosphorylated form (phospho-YAP1) (Fig. 3c, d) as well as their ratio (Fig. 3e) in H69 cells after being stressed with Augmentin at 0.32 and 1 mM for 48 h. TRULI, which is a well-known YAP1 activator, was used as a positive control. Immunocytochemistry confirmed the presence of YAP1-positive cells (Fig. 3f), although no significant differences were observed between Augmentin-treated and non-treated cells (Fig. 3f).

**Fig. 3:**
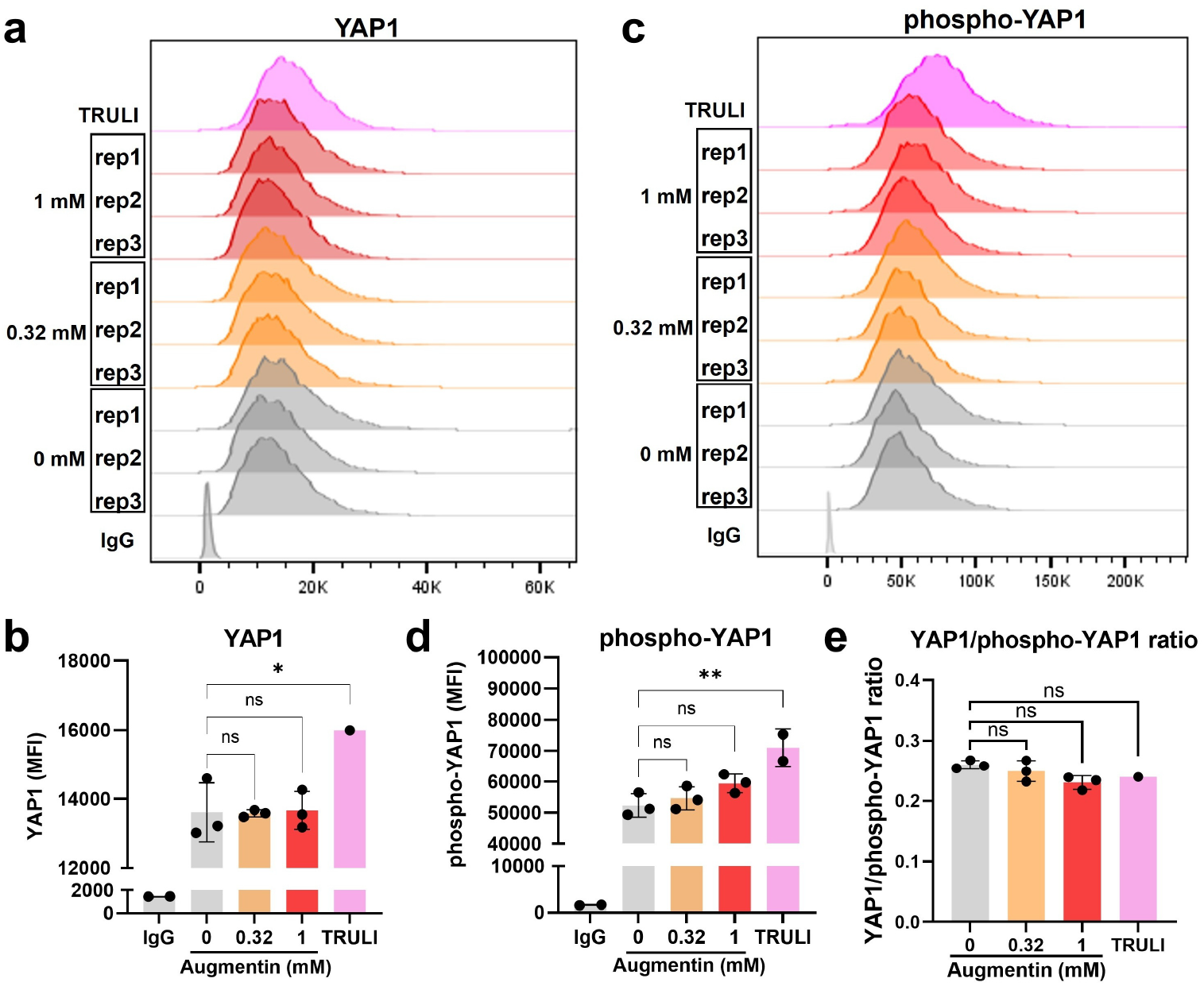
Effects of Augmentin-stress on the cholangiocyte malignancy marker YAP1. **(a-b)** H69 cholangiocytes were stressed with Augmentin at 0.32 and 1 mM for 48 h. After exposure, flow cytometry was performed to evaluate YAP1, a selected cholangiocyte malignancy marker (a), for its comparison based on mean-fluorescence intensities (MFIs, b). TRULI, a reported small-molecule YAP activator, was used as a positive control (10 μM, 48 h). **(c-d)** Flow cytometry was performed to evaluate the phosphorylated form of YAP1 (c) and MFI (d). **(e)** Ratio of YAP1/phospho-YAP1.

The above experiments were carried out in bulk samples, which could miss single-cell events. We therefore examined whether Augmentin stress induces changes in YAP1 at the single EV level. For YAP1 detection in single EVs, we used a more sensitive method based on fluorescence-amplified EV sensing (FLEX) technology(*18*). In FLEX analysis, EVs are captured on periodic gold nanowell arrays in a FLEX sensor chip, and EVs’ weak fluorescence signals are amplified by the underlying plasmonic nanostructures, enabling highly sensitive multichannel marker analysis in single EVs. YAP1 is present inside EVs as an intravesicular marker and therefore requires additional permeabilization. The captured EVs were fixed, permeabilized, and immunostained with blocking (**Fig. 4a**). FLEX offers low EV loss during permeabilization, immunolabeling, and subsequent washing steps, whereas the in-solution staining method used in nanoflow cytometry results in significant sample loss at each step. We also included TSG101 as a positive intravesicular EV marker and CD9 as a positive surface EV marker, along with respective IgG controls (**Fig. 4b**). FLEX analysis showed no significant difference in YAP1 levels in H69-derived EVs between the control and Augmentin-stressed groups (**Figs. 4c-d**). Together, the above results suggest that Augmentin stress under the tested conditions in this study did not upregulate the malignancy marker YAP1, neither at the cellular level nor at the single-EV level.

**Fig. 4:**
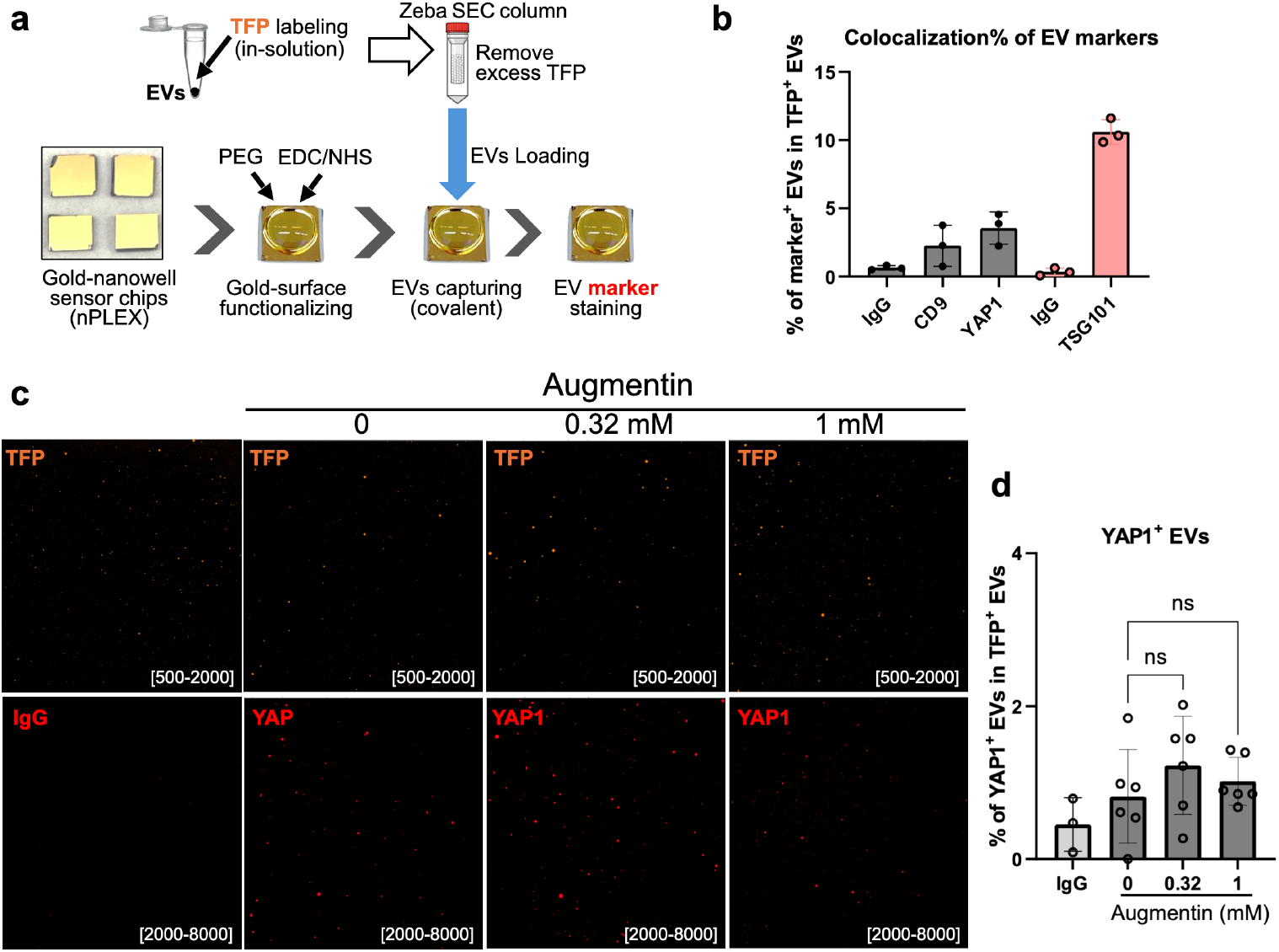
Effects of Augmentin-stress on the cholangiocyte malignancy marker YAP1 by single EV analysis (FLEX). **(a)** Single EV analysis procedure using FLEX sensor chips. The gold surface of the FLEX chip was functionalized by PEG coating. Then, TFP-555-labeled EVs were loaded and captured through covalent binding, followed by immunostaining with fluorophore-conjugated antibodies. Fluorescent images of EVs were acquired and quantified. **(b)** Detection of markers with FLEX using optimized conditions. The percentage of marker-positive EV populations in total TFP-555-positive EVs was calculated (colocalization). **(c-d)** Results of FLEX for the detection of the cholangiocyte malignancy marker YAP1 in Augmentin-stressed cholangiocytes showed no significant change. (c) Representative images of the TFP-555 channel and the YAP1-647 channel. (d) Quantification results of marker colocalization, i.e., the percentage of YAP1-positive EVs in TFP-555-positive EVs.

In addition, we carried out trial screening for other drugs that have been reported to be related to drug-induced liver injury and bile-duct injury, including chlorpromazine, floxuridine, 5-fluorouracil, flucloxacillin and terbinafine. The expression of the malignancy marker YAP1 in the H69 cholangiocyte did not show significant upregulation after treatment with these drugs for 48 h at the tested doses (**Fig. 5**).

**Fig. 5:**
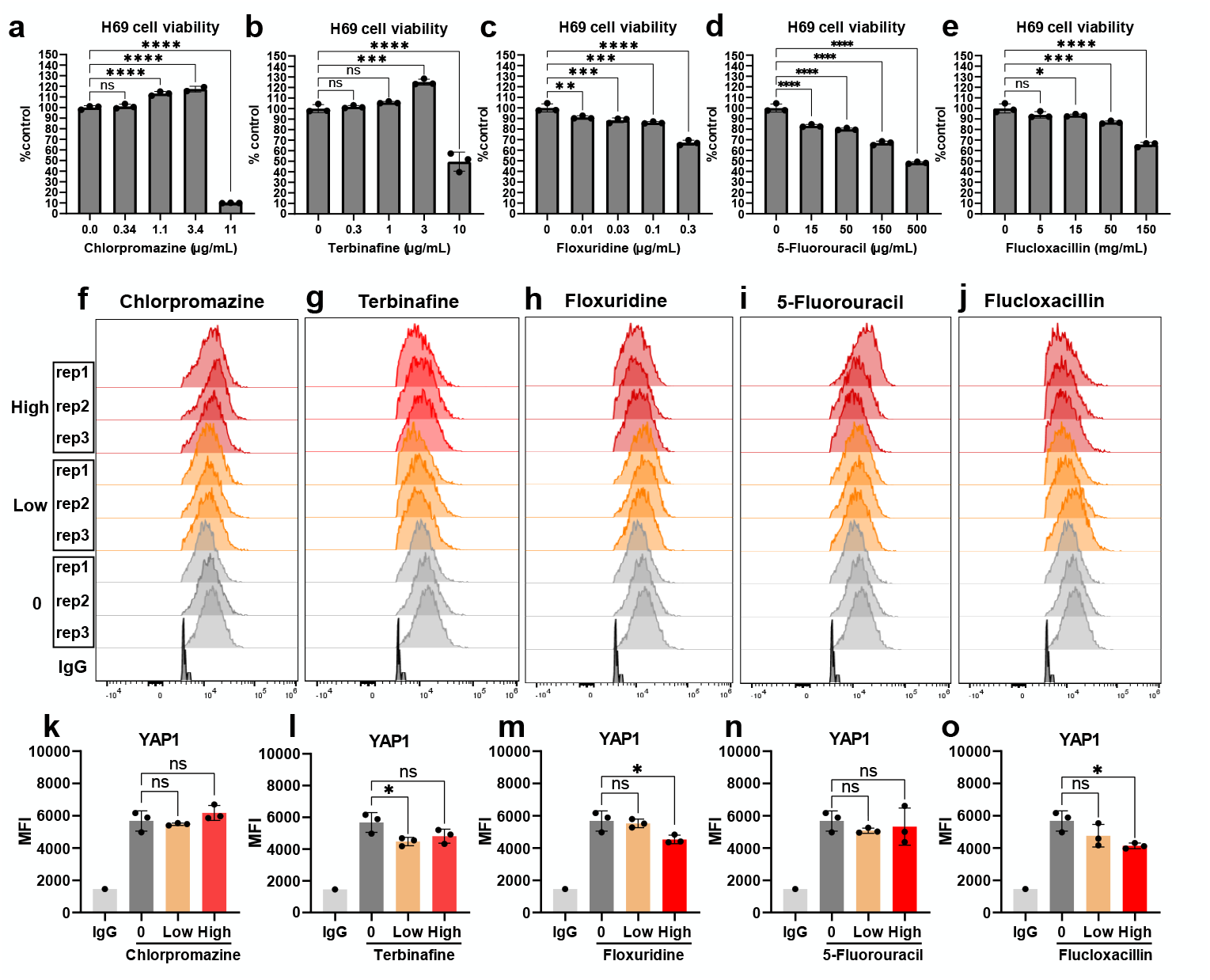
Screening for the malignancy marker YAP1 in H69 cholangiocytes after drug stress. (a-e) Cell viability and (f-o) YAP1 expression in H69 cholangiocytes were measured after treatment with selected drug candidates for 48 h, including chlorpromazine at a low dose of 1.1 µg/mL and a high dose of 3.4 µg/mL (a, f, k); terbinafine at a low dose of 1 µg/mL and a high dose of 3 µg/mL (b, g, l); floxuridine at a low dose of 0.03 µg/mL and a high dose of 0.1 µg/mL (c, h, m); 5-fluorouracil at a low dose of 15 µg/mL and a high dose of 50 µg/mL (d, i, n); and flucloxacillin at a low dose of 15 µg/mL and a high dose of 50 µg/mL (e, j, o).

### Cohort study to explore the association between Augmentin use and CCA risk

To explore the clinical potential association between Augmentin use and CCA risk, we conducted a retrospective cohort study (**Fig. 6a-b**) using the Research Patient Data Registry (RPDR), a centralized longitudinal real-world data repository of clinical electronic health records within the Massachusetts General Brigham enterprise healthcare system spanning 16 member institutions and serving approximately 2.5 million patients annually(*24*). The cohort was constructed using an Active Comparator, New User (ACNU) cohort design(*25*). We selected amoxicillin as the active comparator for Augmentin to reduce bias from indication confounding. Amoxicillin was selected because it has minimal hepatotoxic potential, whereas the liver/bile duct injury associated with Augmentin appears to be primarily attributable to clavulanate rather than amoxicillin(*26*). We first applied preliminary queries using the online RPDR Query tool. Cases with liver/bile duct diseases and carcinoma before the index date were excluded. In the preliminary queries, we recruited 99,587 cases for the control group and 99,014 cases for the Augmentin group. For the acquired raw data, we first conducted quality control and data filtration to obtain valid records containing complete demographics, medication, and diagnosis reports, yielding 58,921 and 66,210 valid cases for the control and Augmentin groups, respectively. Then, a washout period was applied, excluding cases with amoxicillin/Augmentin medication records within 2 years prior to the index date. This resulted in the identification of 35,662 and 38,812 cases as “new-users” for amoxicillin or Augmentin. We further applied an exclusion criterion to remove cases who already had diagnosis records of liver/bile-duct/carcinoma-related diseases before their first use of amoxicillin or Augmentin. This yielded 20,028 and 25,745 cases remaining as the final eligible analytic cohorts for the control and Augmentin groups, respectively. Finally, we followed up the cases in the two groups until the censoring date and identified 16 and 24 CCA cases in the control and the Augmentin group, respectively.

**Fig. 6:**
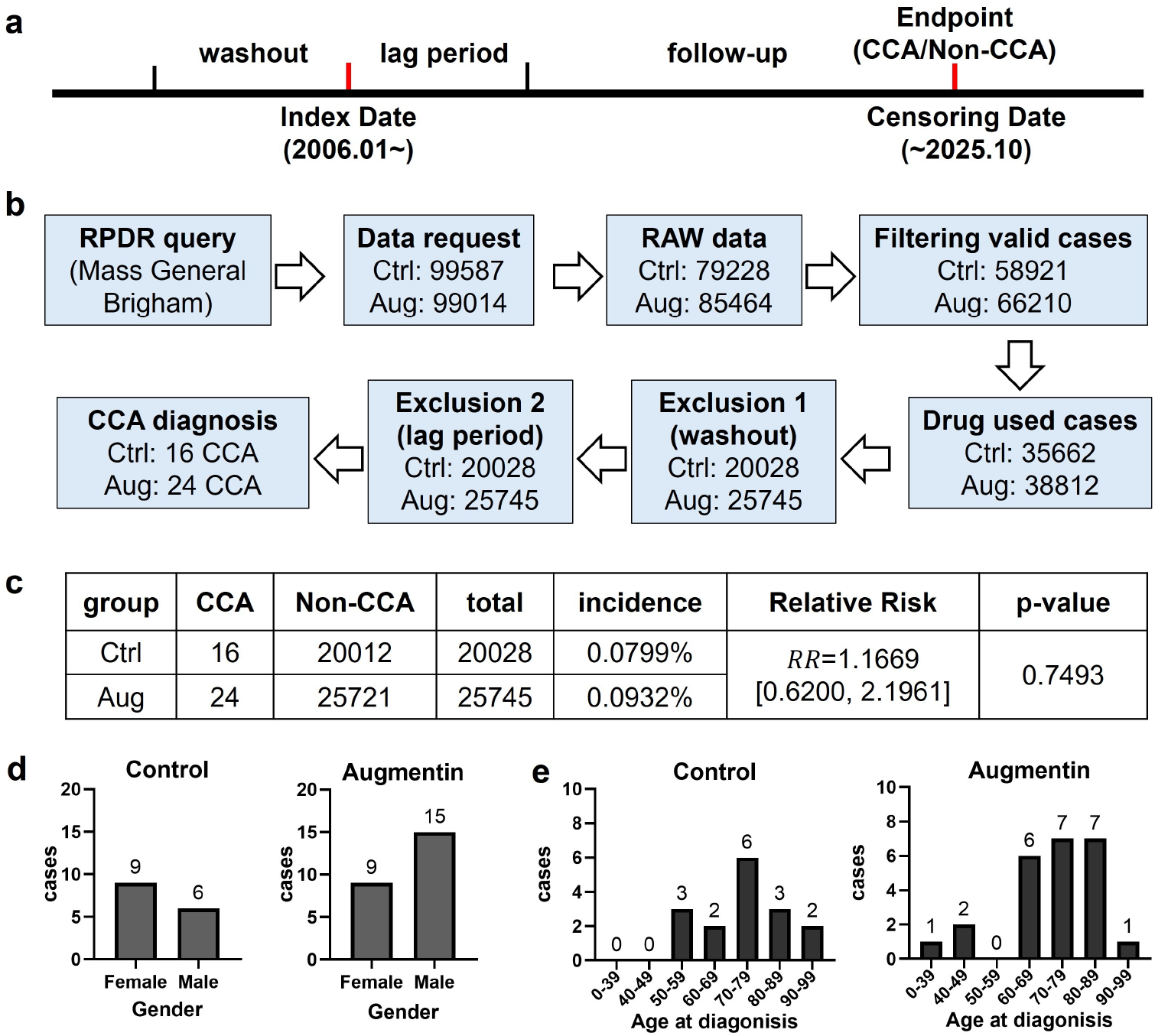
Association between Augmentin use and the risk of CCA: a retrospective cohort study using the RPDR database. (a) Retrospective cohort design. (b) Procedures for processing cohort data. (c) Calculations of CCA incidence in the two groups, relative risk (RR), and p-value. (d) Gender distribution of identified CCA cases in the two groups. (e) Age distribution of identified CCA cases in the two groups.

We further applied a 24-month lag period by excluding patients diagnosed with CCA within 24 months of the first use of amoxicillin or Augmentin (**Fig. 6b**). The results showed that all CCA cases were diagnosed >6 years after the first use of amoxicillin or Augmentin, so no cases were excluded. Finally, the cumulative incidence of CCA was 0.0799% in the amoxicillin group and 0.0932% in the Augmentin group, corresponding to a relative risk (RR) of 1.1669 (95% CI, 0.6200-2.1961). Results of statistical calculation showed that although the RR point estimate was above 1.0, suggesting a numerically higher CCA incidence in the Augmentin group, the association did not reach statistical significance (P=0.7493, Fisher’s exact test) (**Fig. 6c**). Among the identified CCA cases, the control group included 9 female and 6 male patients, whereas the Augmentin group included 9 female and 15 male patients, indicating a male predominance in the Augmentin group (**Fig. 6d**). No CCA cases younger than 50 years were observed in the control group, whereas 3 such cases were identified in the Augmentin group (**Fig. 6e**). Overall, CCA cases in the control group were mainly concentrated in the 70–79 year group, while cases in the Augmentin group were more broadly distributed, with the highest numbers in the 70–79 and 80–89 year groups (**Fig. 6e**).

## Discussion

Cholangiocarcinoma development has been linked to carcinogen exposure, including a long list of toxins and therapeutically used drugs. Among the latter, it has been suggested that certain antibiotics, including Augmentin, may cause CCA. We hypothesized that, if such a link exists, it could be detected via ultra-sensitive vesicle analysis of cholangiocytes exposed to the drug. We therefore developed a system that enabled controlled exposure of cholangiocytes to drugs, the harvesting of EVs produced, and their analysis at the single-EV level. The single-EV analysis technology had been described before(*18, 27, 28*) but had not been adapted to YAP1/pYAP1 measurements in EV. We show that the system is well-suited for stress analysis of cholangiocytes and that positive controls for both intravesicular and surface markers (TSG101 and CD9, respectively) are indeed positive.

We tested the potential for malignant transformation of several drugs, but focused primarily on Augmentin since it is so widely used. Augmentin is a commonly prescribed antibiotic that combines amoxicillin and clavulanic acid. Amoxicillin works by interfering with the formation of bacterial cell walls. Clavulanic acid blocks beta-lactamases, protecting amoxicillin from being broken down. It has been suggested that clavulanic acid is the culprit in cholangiocyte toxicity, and therefore, it provided a model system in which amoxicillin alone could serve as the control. In essence, our results show no direct experimental or clinical evidence that Augmentin causes malignant transformation of cholangiocytes at the doses tested.

The current study, however, presents several limitations. First, the in vitro model primarily examined cellular responses at a single time point (48 h) while the cohort study included a full clinical regimen. Future studies incorporating repeated exposures to achieve prolonged or cumulative exposure durations would provide additional insight into long term exposure. Second, given the critical roles of immune and inflammatory responses in Augmentin-induced liver injury(*29, 30*), further investigations into the interactions between cholangiocytes and other relevant cell types could be warranted. For example, co-culture systems integrating macrophages, hepatocytes, and cholangiocytes may offer a more physiologically relevant model, but this is generally difficult to achieve as these models rarely reflect the in vivo dynamic interaction of immune cell subsets and spatial arrangements. Third, in the cohort analysis, the number of observed CCA events was small given the low incidence of CCA despite the fact that we started with nearly 200,000 patients who had received antibiotics in question. Further large-cohort meta-analysis in countries with a higher incidence rate may provide additional insight

Despite these limitations, this study provides valuable information and establishes a foundational framework, including both the stress model and the analytical approach, which may serve as a reference for the development of more complex experimental systems in the future. In particular, our work highlights the promising prospects of applying the single-EV analysis approach in the fields of pharmaceutical research and characterizing molecular responses to therapy (e.g., adverse effect screening, postmarketing pharmacovigilance), as well as in toxicology fields, including applications in biomarker development for toxicity evaluation, elucidation of molecular mechanisms, and identification of potential therapeutic targets.

## Materials and Methods

### Reagents and preparation of exposure solutions

Reagents for exposure experiments were all commercially purchased, including Augmentin (amoxicillin trihydrate/potassium clavulanate; Cat. SMB00607, Sigma-Aldrich); Floxuridine (Cat. HY-B0097, MedChem Express); 5-Fluorouracil (Cat. HY-90006, MedChem Express); Flucloxacillin sodium (Cat. HY-A0246A, MedChem Express); Terbinafine (Cat. HY-17395A, MedChem Express); Chlorpromazine hydrochloride (Cat. HY-B0407A, MedChem Express); and TRULI (Cat. SML3636, Sigma-Aldrich). Stock solutions of the above reagents were prepared by dissolving them in dimethyl sulfoxide (DMSO; Cat. D8418, Sigma-Aldrich). Aliquots of the stock solutions were stored at -80 °C until use. For cell treatment, the stock solutions were thawed and diluted immediately before use in culture medium (DMEM/DF-12 at 1:1 in volume) supplemented with 2% exosome-depleted FBS (Cat. A2720801, Thermo Fisher Scientific). The final DMSO concentration in the exposure solutions was adjusted to 0.1%. To initiate the exposure, the cells were washed twice with PBS, and the prepared exposure solution was added. The cells were then incubated at 37 °C and 5% CO_2_ for the indicated time.

### Cell culture

The H69 cell line, a human normal cholangiocyte cell line, was generously provided by Prof. Yangmi Kim and Prof. Seon Mee Park at Chungbuk National University College of Medicine(*18*). H69 cells were maintained in a 1:1 (v/v) mixture of culture medium DMEM (Cat. 10-013-CV, Corning) and DMEM/F-12 (DF12, Cat. 11320033, ThermoFisher Scientific), supplemented with 10% heat-inactivated fetal bovine serum (FBS; Cat. A5256801, ThermoFisher Scientific), 0.025 mg/ml adenine (Cat. A8625, Sigma-Aldrich, St. Louis, MO, USA), 0.005 mg/ml insulin (Cat. 12585014, Gibco Invitrogen), 0.001 mg/ ml epinephrine (Cat. E4250, Sigma-Aldrich), 13.6 ng/ml 3,3′,5-Triiodo-L-thyronine (T3) (Cat. T6397, Sigma-Aldrich), 0.0083 mg/ml holo-transferrin (Cat. T0665, Sigma-Aldrich), 620 ng/ml hydrocortisone (Cat. H0888, Sigma-Aldrich) and 10 ng/ml epidermal growth factor (EGF; Cat. 236-EG-200, R&D Systems) at 37 °C in 5% CO_2_. The culture medium was freshly prepared and used within one month. For cell passage, the subconfluent cells were washed with PBS (Cat. 21040CV, Corning) and detached with 0.05% Trypsin-0.53 mM EDTA (Cat. 25-052-CI, Corning). The cells were regularly tested for mycoplasma using the Universal Mycoplasma Detection Kit (ATCC), and no contamination was detected. H69 cells at passage numbers 8-15 were used for experiments.

### Cell viability assay

Cell viability assay was performed using Cell Counting Kit-8 (CCK-8) (Cat. HY-K0301, Med Chem Express), following the manufacturer’s instructions. Briefly, cells were seeded into a 96-well plate at a density of 1×10^4^ cells per well in 100 μL and cultured overnight. Then the cells were washed with PBS, followed by drug treatments at the indicated doses and for the indicated time. At the end of exposure, 10 μL of CCK-8 reagent was added directly to each well and incubated for 1.5 h at 37 °C and 5% CO_2_. The absorbance was then measured at 450 nm with a Spark Multimode Microplate Reader (TECAN, Männedorf, Canton of Zürich, Switzerland). Finally, cell viability was calculated and expressed as the percentage relative to the control group.

### Flow cytometry for cells

After treatment, cells were washed with PBS and detached using 0.05% Trypsin-0.53 mM EDTA for 5 min, followed by neutralization with culture medium containing 10% FBS. Then, after washing with PBS twice, cells were stained with the Zombie Violet Fixable Viability Kit (Cat. 423113, BioLegend) at a 1:100 dilution in PBS for 15 min at room temperature (RT), and washed with 0.1% BSA (Cat. 10735086001, Roche Diagnostics GmbH, Mannheim, Germany) in PBS. The cells were then fixed in 4% paraformaldehyde (PFA; Cat. 15710, Electron Microscopy Sciences) for 30 min at RT, followed by membrane permeabilization with Perm Reagent (True-Nuclear Perm reagent, Cat. 73162, BioLegend) for a total of 15 min at RT. The cells were blocked with 2% BSA for 1 h at RT, followed by antibody staining. For YAP1 staining, the cells were incubated with the primary antibody (mouse anti-human YAP1, Alexa Fluor 647-conjugated, sc-376830, Santa Cruz) at a 1:100 dilution in Perm reagent for 1 h at RT. For phospho-YAP1, the cells were incubated with primary antibody (rabbit anti-human phospho-YAP1, unconjugated, Cat. 80694-2-RR, ProteinTECH) at 1:500 dilution in Perm Reagent for 1 h at RT, followed by incubation with secondary antibody (Donkey anti-rabbit IgG, Alexa Fluor 647 conjugated, Cat. 406414, BioLegend) at 1:400 dilution in Perm Reagent for 30 min at RT. Isotype controls of primary antibodies were used, including Mouse IgG1 kappa Isotype Control-AF647 conjugated (Cat. 51-4714-81, ThermoFisher Scientific) and Rabbit IgG Isotype Control (Cat. 14-4616-82, Invitrogen). Finally, the cells were washed with Perm Reagent to remove excess antibodies and resuspended in 0.5% BSA in PBS. The cells were then analyzed using a flow cytometer (Attune NxT Flow Cytometer, Thermo Fisher Scientific). The data were acquired with Attune Cytometric software (version 5.3.0) and analyzed with FlowJo(version 10.9.0, TreeStar).

### Immunocytochemistry

For immunocytochemistry experiments, a glass-bottom chamber (Cat. 154534, Nunc Lab-Tek II Chamber Slide System, ThermoFisher Scientific) was used for cell seeding and drug treatment. After treatment, the cells were washed with PBS and fixed in 4% PFA for 30 min at RT, followed by membrane permeabilization with Perm Reagent (True-Nuclear Perm reagent, Cat. 73162, BioLegend) for 15 min at RT. Then the cells were blocked with 2% BSA for 1 h at RT, followed by incubation with the primary antibody (mouse anti-human YAP1, Alexa Fluor 647-conjugated, sc-376830, Santa Cruz) at a 1:100 dilution in Perm Reagent for 1 h at RT. Finally, the cells were incubated with DAPI (1 μg/mL, Cat. D9542, Sigma-Aldrich) for nuclear staining and mounted with ProLong™ Glass Antifade Mountant (Cat. P36982, ThermoFisher Scientific). A glass coverslip was then applied, and the cells were observed using a BX63 Upright Epifluorescence Microscope (Olympus).

### EV isolation

Size-exclusion chromatography (SEC) was used to isolate EVs from the conditioned medium of drug-treated cells. Briefly, cells were grown to 70% confluence, washed twice with 0.22 μm-filtered PBS, and the medium was replaced with exposure solution prepared in medium supplemented with 2% exosome-depleted FBS (Thermo Fisher Scientific, Cat. A2720801) for the indicated time, followed by collection of the conditioned medium and EV isolation. The conditioned medium was centrifuged at 300 g for 10 min and 2000 g for 20 min at 4 °C, then filtered through a 0.8 μm filter to remove cell debris. The conditioned medium was concentrated with Centricon Plus-70 Centrifugal Filter (MWCO 100 kDa, Cat. UFC9100, Sigma-Aldrich) by centrifuging at 3500 g for 30 min at 4 °C. Then EVs were purified from the concentrated medium by passing through a SEC column (Econo-Pac® Chromatography Columns, Cat. 7321010, Bio-Rad), which was pre-filled with Sepharose CL-4B resin (Cat. 17015001, Cytiva).

The 4^th^ and 5^th^ fractions were used for EV isolation. The collected EVs were further concentrated using an Amicon Ultra-2 Centrifugal Filter (MWCO 100 kDa, Cat. UFC210024, Sigma-Aldrich) by centrifuging at 3500 g for 30 min at 4 °C. Finally, the concentrated EVs were aliquoted into several tubes and stored in a −80 °C deep freezer until analysis.

### Pan-EV staining with fluorescent TFP-555 dye

To separate EVs from non-EV particles, pan-EV staining was performed using an optimized ubiquitous EV labeling approach, with fluorescent tetrafluorophenyl (TFP) labeling primary amines on EV-surface proteins. Previous work from our group has reported that this amine-labeling approach is superior to other lipid staining methods and showed no anticipated bias towards different-sized EVs(*18, 28, 31*). To prepare the fluorescent TFP dye, 27.5 mM Azido-dPEG®_12_-TFP ester (Cat. QBD-10569-100, Vector Laboratories) was mixed with 25 mM AZDye 555 DBCO (Cat. CCT-1290-1, Vector Laboratories) at 1:1 (v/v) and incubated for 2 h at RT. Then the obtained TFP-555 dye was aliquoted and stored at -80 °C until use. For pan-EV staining, 1 μL of EVs was mixed with 2 μL PBS, 2 μL 100 mM sodium bicarbonate (Cat. 25080094, ThermoFisher Scientific), and 0.2 μL the TFP-555 dye to a total volume of 5.2 μL and incubated in the dark at RT for 1 h on a HulaMixer Rotating Mixer (ThermoFisher Scientific). Then the excess TFP-555 dye was removed by passing it through a Zeba™ Spin Desalting Column (MWCO 40K, Cat. A57758, Thermo Fisher Scientific).

### Single EV analysis by nanoflow cytometry

Total EV count and EV markers were examined by nanoflow cytometry (CytoFLEX Nano, Beckman Coulter), an adapted flow cytometry method for nanoscale particles. For EV count measurement, isolated EV samples were diluted in PBS and directly analyzed by nanoflow cytometry. To measure EV surface markers (e.g., CD9, CD63, CD81), EVs were first labeled with the fluorescent TFP-555 dye as described above. Then the EVs were blocked by adding 25 μL of 2% BSA and incubated for 1 h in the dark at RT on a HulaMixer. Then the EVs were incubated with the indicated antibodies for 1 h in the dark at RT on a HulaMixer. Finally, after antibody incubation, the EVs were purified by removing excess antibodies using a SEC column (Poly-Prep Chromatography Columns, Cat. 7311550, Bio-Rad) packed with Sepharose CL-4B resin. The acquired EVs were analyzed by nano flow cytometry (CytoFLEX Nano, Beckman). For EV marker quantification, the percentage of marker-positive EV populations among total TFP-555-positive EVs (colocalization) was calculated.

The primary antibodies used are as follows: anti-CD9 (Cat. 312102, BioLegend), anti-CD63 (Cat. 353039, BioLegend), and anti-CD81 (Cat. sc-166029, Santa Cruz). Isotype controls of primary antibodies were used, including Mouse IgG1 kappa Isotype Control-AF647 conjugated (Cat. 51-4714-81, ThermoFisher Scientific). For unconjugated primary antibodies, ReadyLabel Antibody Labeling Kit-Alexa Fluor 647 (Cat. R10710, ThermoFisher Scientific) was used to label the antibodies following the manufacturer’s instructions.

### Single EV analysis by FLEX assay

The fluorescence-amplified extracellular vesicle sensing (FLEX) assay was performed using a nanoplasmonic sensor chip as described in a previous publication(*18*). The FLEX chip is fabricated with periodic gold nanowell arrays on a gold surface to enhance fluorescent signals via long-range surface plasmon resonance. Briefly, the FLEX chip was first cleaned with acetone, IPA, and Mili-Q water. Then, the gold surface was functionalized with a PEG coating using a mixture of 2.5 mM SH-PEG-COOH (PG2-CATH-1k, Nanocs) and 7.5 mM mPEG-SH (PG1-TH-350, Nanocs) at RT overnight, which resulted in the maximum EV capture efficiency(*32*). After washing with PBS, the gold surface was incubated in a freshly prepared mixture containing 0.4 M EDC (Cat. 22980, Thermo Fisher Scientific) and 0.1 M sulfo-NHS (Cat. PG82071, Thermo Fisher Scientific) dissolved in 0.5 M MES (Cat. J62574.AE, Thermo Fisher Scientific) at pH 6.0 for 7 min. After gently washing with PBS, EVs were loaded onto the chip surface and incubated for 30 min at RT. After EV capture, EVs were fixed with 4% PFA for 10 min, followed by incubation with 1M pH8.0 ethanolamine (Cat. E9508, Sigma-Aldrich) for 20 min and permeabilization with 0.01% Triton X-100 (Cat. X100-100ML, Sigma-Aldrich) for 10min. After blocking with 2% BSA for 20 min, the captured EVs were then immunofluorescently labeled with conjugated primary antibodies for 60 min (e.g., mouse anti-human YAP1, AF647-conjugated, Cat. sc-376830, Santa Cruz), or with unconjugated primary antibodies (e.g., mouse anti-human TSG101/Cat. 67381-1, ProteinTech, mouse anti-human CD9/Cat. 312102, BioLegend) followed by incubation with a conjugated secondary antibody (AlexaFluor 647 anti-mouse, Cat. A-21236, Thermo Fisher Scientific) for 30 min. Isotype controls of primary antibodies were used, including Mouse IgG1 kappa Isotype Control-AF647 conjugated (Cat. 51-4714-81, ThermoFisher Scientific) and unconjugated Purified Mouse IgG1, κ Isotype Ctrl Antibody (Cat. 400101, BioLegend). Antibodies were diluted in 1% BSA solution, and staining was performed with gentle agitation. Finally, after antibody incubation, the chips were mounted with ProLong™ Glass Antifade Mountant (Cat. P36982, Thermo Fisher Scientific) and covered with a glass coverslip. Fluorescence images of EVs were acquired with an upright automated epifluorescence microscope (Zeiss AX10, Axio Imager M2) equipped with a CMOS camera (C15440-20UP, Hamamatsu, Japan).

### Image processing and FLEX result quantification

Images acquired from the FLEX assay were analyzed using ImageJ (version 1.54p, NIH) and custom-built Python code(*18*). Briefly, background signals were subtracted using a rolling ball algorithm (radius = 20). Then the ComDet plugin in ImageJ was used to detect EV locations in the AF555 channel (TFP) and the AF647 channel (marker). Averaged fluorescence intensities were calculated from a 3 × 3 fixed pixel window. The obtained data were analyzed using custom-built Python code to calculate the colocalization percentage (the ratio of marker-positive EVs to TFP-positive EVs).

### Retrospective cohort with RPDR database

The retrospective cohort study was conducted using the Research Patient Data Registry (RPDR) under an institutionally approved IRB Protocol 2019P003441. The cohort was constructed with the Active Comparator, New User (ACNU) cohort design. Amoxicillin was selected as the active comparator for Augmentin to reduce bias from indication confounding.

First, preliminary queries were carried out using the online RPDR Query tool. An Index Date was set (January 2006) for cohort accrual. Cases with records of amoxicillin/ Augmentin use after the Index Date were recruited to the cohort. Cases with liver/bile duct diseases and carcinoma before the Index Date were excluded. We recruited 99,587 cases for the control group and 99,014 cases for the Augmentin group. Then, raw data requests were submitted to RPDR and downloaded under the approved IRB protocol. The acquired raw data spanned several domains, including demography, medication, and diagnosis records. We first conducted quality control and data filtering to retain records with complete demographic, medication, and diagnosis data. Then, a washout step was applied, excluding cases with amoxicillin/Augmentin medication records within 2 years before the index date. We further applied an exclusion criterion to remove cases that already had diagnosis records of liver/bile-duct/carcinoma-related diseases before their first use of amoxicillin or Augmentin. Finally, we followed up the cases in the two groups until the Censoring date (October 2025) to identify cases with DIA records for CCA. We further applied a 24-month lag period by excluding patients diagnosed with CCA within 24 months after the first use of amoxicillin or Augmentin. Finally, the cumulative incidence of CCA in the two groups and the relative risk (RR) were calculated. Statistical significance was assessed using Fisher’s exact test.

### Statistical analysis

All statistical data analyses were performed using GraphPad Prism 9 (GraphPad Software Inc., San Diego, CA, USA), and the results are expressed as mean ± standard deviation. For normally-distributed datasets, we used 2-tailed Student’s t-test and one-way ANOVA followed by Dunnett’s multiple comparison test. When variables were not normally distributed, we performed non-parametric Mann-Whitney or Kruskal-Wallis tests. p-values> 0.05 were considered not significant (n.s.), p-values < 0.05 were considered significant.

## Acknowledgments

We thank Dr. Claudio Vinegoni for help with microscopy, Drs. Christopher Garris and Sepideh Parvanian for help with flow cytometry, and Drs. Jihye Hong, Shingo Noguchi, Yen Nguyen, Juhyun Oh, and Grant Simpson for many helpful discussions. We thank all members of Weissleder and Im Labs for their kind support and help. C.Z. was supported in part by the Overseas Research Fellow Program of Tokyo University of Science. This work was supported in part by the following NIH grants: T32 5T32CA079443-24 (S.A.W.), R33CA277820, R01 CA293063 (R.W.), R01GM138778 (H.I.), and R33CA281794 (H.I.). R.G. was supported by the NCI grant 5R25CA174650 through the CanCURE program at Northeastern University. K.L and J.H.J. were supported by the National Research Foundation of Korea (NRF) grant (RS-2024-00335625).

## Contributions

Conceptualization: CZ, HI, RW

Data curation: CZ, HI, RW

Formal analysis: CZ

Methodology: CZ, KL, SAW, RD, MHJ, RG, JHJ, SI, HI, RW

Software: CZ, KL

Cohort study: CZ, RD, SI, RW

Supervision: HI, RW

Visualization: CZ, RW

Writing - original draft: CZ, RW

Writing - review and editing: RW and all coauthors

Funding acquisition: RW

Project administration: RW

Resources: RW

